# Multimodal profiling of proinflammatory protease activity identifies caspase-1 as a target for lung cancer interception

**DOI:** 10.1101/2025.07.01.662195

**Authors:** Cathy S. Wang, Qian Zhong, Shih-Ting Wang, Carmen Martin-Alonso, Sofia Neaher, Sahil Patel, Tiziana Parisi, Jesse D. Kirkpatrick, Lecia V. Sequist, Tyler E. Jacks, Sangeeta N. Bhatia

## Abstract

The systemic inhibition of IL-1b, a key mediator of pulmonary inflammation, has been shown to reduce the incidence of lung cancer in patients in years following treatment, but knowledge gaps surrounding its activation and role in the tumor microenvironment are hindering approaches for cancer interception. We developed a suite of activity-based technologies to probe inflammation in early lung cancer and identified a translational target candidate. We designed probes sensitive to various IL-1b-activating proteases and applied them to a murine model of inflammatory lung cancer, *Kras/Trp53*-mutant with SIINFEKL expression (KPS). Our nanosensors revealed reduced cleavage of a caspase-1 reporter in the lungs of KPS mice treated with IL-1b antibody, as well as elevated caspase-1 expression and activity in naïve tumor tissue sections, highlighting the importance of caspase-1 processing of IL-1b during cancer development. We conducted a pre-clinical trial of a novel combination intervention by administering both IL-1b blockade and caspase-1 inhibition shortly after tumor induction. Following treatment, we observed significant reduction in lung cancer formation, including complete ablation of tumor incidence in nearly 20% of KPS mice. Our approach to understand the interplay of protease activity and cytokine activation supports development of new strategies to mitigate inflammation and intercept lung cancer progression.

## Introduction

Lung cancer is the second most common cancer globally, as well as the leading cause of cancer death in both men and women in the United States [1]. Despite advances in chemo- and immunotherapies, 5-year survival rates are still low due to late diagnosis. Cancer interception emerges as a crucial approach to detect and prevent cancer development. A promising indication for lung cancer interception was revealed by the CANTOS trial in 2017, in which systemic administration of IL-1b antibody led to a significant reduction in lung cancer incidence and mortality in patients years later [2]. However, when IL-1b blockade was applied to patients with late-stage non-small cell lung cancer (NSCLC) in subsequent CANOPY trials, very little improvement in survival was observed [3]. This finding suggests that IL-1b inhibition might offer an opportunity for cancer interception with sufficiently early anti-inflammatory intervention. This possibility has the potential to drastically alter lung cancer outcomes, but to reach this goal, a greater understanding of the factors that influence IL-1b-associated inflammation pathways in early tumorigenesis is needed.

IL-1b is a particularly appealing pathway to target, due to its pro-inflammatory function in both acute and chronic inflammation, which have been shown to lead directly to lung cancer. IL-1b transcription is triggered in response to Toll-like receptor engagement, and IL-1b’s binding to IL-1Rs on a wide variety of cells activate MAPK and NF-kB pathways, leading to inflammatory cytokine induction and immune cell recruitment [4, 5]. IL-1b is upregulated in tumor and adjacent lung tissue in NSCLC, and high serum protein levels in lung cancer patients have been linked to poor survival [6, 7]. Hill et al showed that IL-1b antibody blockade in EGFR-mutant mice during exposure to particulate matter reduced lung adenocarcinoma formation [8]. Other studies have observed that IL-1b can lead to activation of immunosuppressive cells including M2 macrophages and myeloid-derived stem cells, as well as increased epithelial-to-mesenchymal transition and invasiveness of cancer cells [9, 10]. IL-1b can act via two pathways: inflammasome-dependent and -independent. In these two pathways, IL-1b is produced as an inactive pro-form by different cell types and can be activated by multiple proteases [11]. Thus, IL-1b’s proinflammatory functions, and more broadly, chronic inflammation-induced lung cancer, depend on these immune-related IL-1b-cleaving proteases [12]. However, there is a lack of clarity for which proteases participate and how cleavage-mediated IL-1b activation influences local cell populations towards tumorigenesis. Uncovering the role of these proteases in chronic inflammation-induced lung cancer will lead to new approaches for targeted perturbation and resulting cancer interception.

In this study, we utilized an autochthonous murine model of lung adenocarcinoma (LUAD), Kras/Trp53-mutant mice with SIINFEKL expression (KPS) that creates an inflammatory environment eliciting tumor growth starting at 4-6 weeks post-induction [13]. To investigate the involvement and impact of protease activity in the progression of disease, we leveraged different proteases’ ability to recognize and cleave unique amino acid sequence motifs and designed a library of 124 Fluorescence Resonance Energy Transfer (FRET) substrates that included motifs for interleukin-processing proteases. After down-selection screening with recombinant proteases, 20 substrates were prioritized to create *in vivo* activity-based nanosensors (ABNs). ABNs, which have previously been used to track protease activity changes for disease diagnosis, can be used to profile which proteases contribute to disease progression and measure changes in the presence or absence of treatments [14-16]. This panel of ABNs monitored KPS tumor response to weekly administration of IL-1b and PD-1 blockade. With this treatment, our studies revealed a reduction in caspase-1 activity, known for its role in processing IL-1b [17]. We found robust localization of caspase-1 activity to developing lung tumor nodules by applying a third peptide-based technology, activity-based zymography probe, that fluorescently “paints” tissues that exhibit protease activity [18]. Based on these findings, we hypothesized that caspase-1 activity contributes to the production of active IL-1b in LUAD development. As such, using a clinically safe small molecule inhibitor of caspase-1, we studied the effect of caspase-1 inhibition, alone and in combination with IL-1b blockade, *in vivo*. Caspase-1 inhibition resulted in a significant reduction in cancer progression in KPS mice, and a repeatable fraction of combination-treated mice had no detectable tumors. Collectively, we demonstrate how activity-based nanoscale technologies help to elucidate the inflammatory microenvironment in early cancer development and propose a novel therapeutic target for lung cancer interception.

## Results

### Peptide substrates were nominated for cleavage by IL-1b-activating proteases

Protease dysregulation has been known to play a role in oncogenic processes such as promoting growth, angiogenesis, and inflammation [19]. Protease activity can be detected *in vitro* and *in vivo* through measuring fluorescent and mass-encoded readouts from cleavage of amino acid sequence substrates that are designed to be recognized by target proteases [14, 15, 20]. As the cleavage and resulting activation of IL-1b in inflammation is also governed by proteases, we sought to monitor these upstream activators to identify potential changes that contribute to inflammation-associated lung cancer (Fig. 1A). We designed a library of 124 substrates consisting of 8-13 amino acid sequences with motifs that would be recognized by proteases that cleave IL-1a, IL-1b, IL-18, and IL-33 (Fig. 1B, Table S1) [19, 21, 22]. Sequences that were successfully synthesized with a fluorophore-quencher pair were used in several rounds of FRET screening. Fluorescence was only detected when cleavage of the substrates separated the quencher from the fluorophore. We assayed FRET substrates against recombinant proteases of interest to determine the specificity and efficacy with which each protease cleaves a given substrate (S1). We found various substrates that were recognized and cleaved effectively by at least one of the proteases. Due to the known promiscuity of proteases, some substrates such as #28, #29, and #70 were cleaved efficiently by multiple proteases of the same class [20].

**Figure 1.**
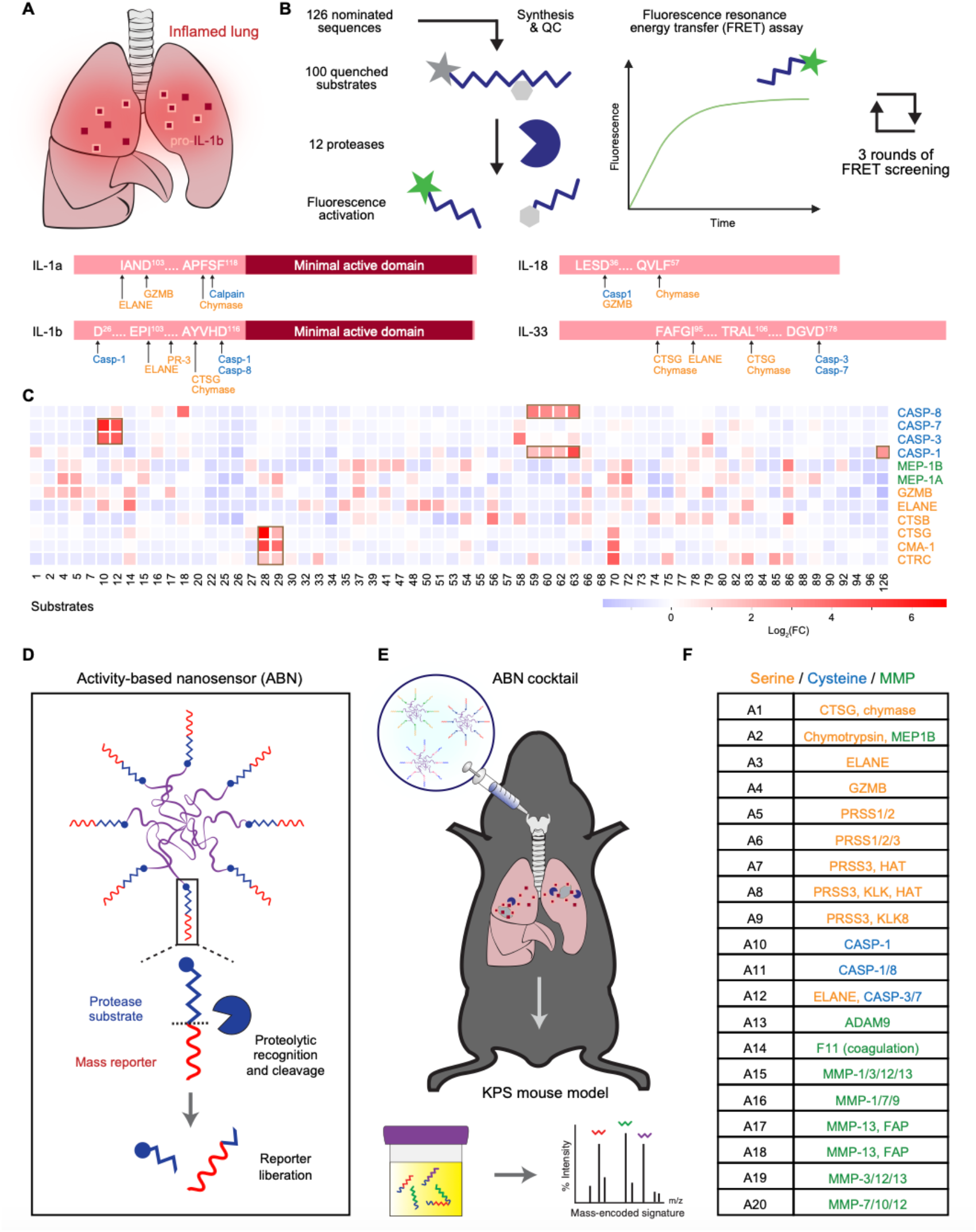
Interleukin-associated protease substrates were designed and screened to develop an in vivo sensor panel. (**A**) Schematic of inflamed lung with high expression of IL-1b in pro-form and cleavage sites of associated interleukins [16]. (**B**) Schematic of FRET protease substrate screening. 100 amino acid sequences with fluorophore-quencher pairs were designed or nominated from previous lung cancer diagnostic panels and incubated with 14 proteases across three rounds of FRET cleavage assays, where fluorescence increases over time with cleavage. (**C**) Heatmap of z-scored cleavage fold change from a representative FRET screen. Yellow boxes indicate examples of strong cleavage trends for select probes. (D) Schematic of activity-based nanosensor (ABN) cleavage by local proteases *in vivo*. (**E**) Intratracheal administration and mass spectrometry readout shows cleaved ABN reporters in urine from mice. (**F**) List of ABN probes and corresponding target protease selected for *in vivo* administration.

For dynamic and longitudinal *in vivo* sensing of these proteases, we converted substrates that were efficiently cleaved by our proteases of interest into linkers for activity-based nanosensors (ABNs). To form these nanoparticles, PEG scaffolds are conjugated with unique mass-encoded tags via different cleavable substrate linkers (Fig. 1D) [23]. Upon intratracheal delivery of ABNs, local proteases in the lung can recognize and cleave specific amino acid sequence motifs, which releases the mass reporters into circulation for renal concentration and urinary clearance. Mass spectrometry of the collected urine samples reveals the amount and identity of the reporter tags, and by proxy, the protease-cleaved substrates (Fig. 1E). Our multiplexed panel of ABNs consisted of 20 substrates of previously validated lung cancer targets, as well as hits from screening recombinant IL-1b-activating proteases, designed to detect changes in protease activity in the lung with IL-1b-associated intervention (Fig. 1F, Table S2) [14].

### IL-1b blockade in inflammatory lung cancer model results in differential protease cleavage

The ABNs were applied to the LUAD KPS murine model, where lung tumors were induced in Kras/Trp53-mutant mice by intratracheal injection of lentivirus encoding for both Cre recombinase and SIINFEKL. The expression of the SIINFEKL antigen in KPS mice creates a more immunogenic lung environment in which the tumors grow, as shown through early IL-1b expression in tumor nodules starting from six weeks (S2) [13, 24]. In congruence with prior studies, we adopted a dosing regimen of weekly administration with either control IgG antibody, IL-1b antibody, PD-1 antibody, or combined IL-1b and PD-1 antibodies to KPS mice, starting at five weeks after tumor inoculation (Fig. 2A) [13]. We observed that introducing IL-1b blockade at these early stages reduced tumor formation, consistent with previous observations that IL-1b plays a role in tumor development in this LUAD model (Fig. 2B) [13]. We then wanted to characterize the pattern of protease activity in the early inflammatory environment and detect when and how activation of the IL-1b pathway might play a role in cancer progression.

**Figure 2.**
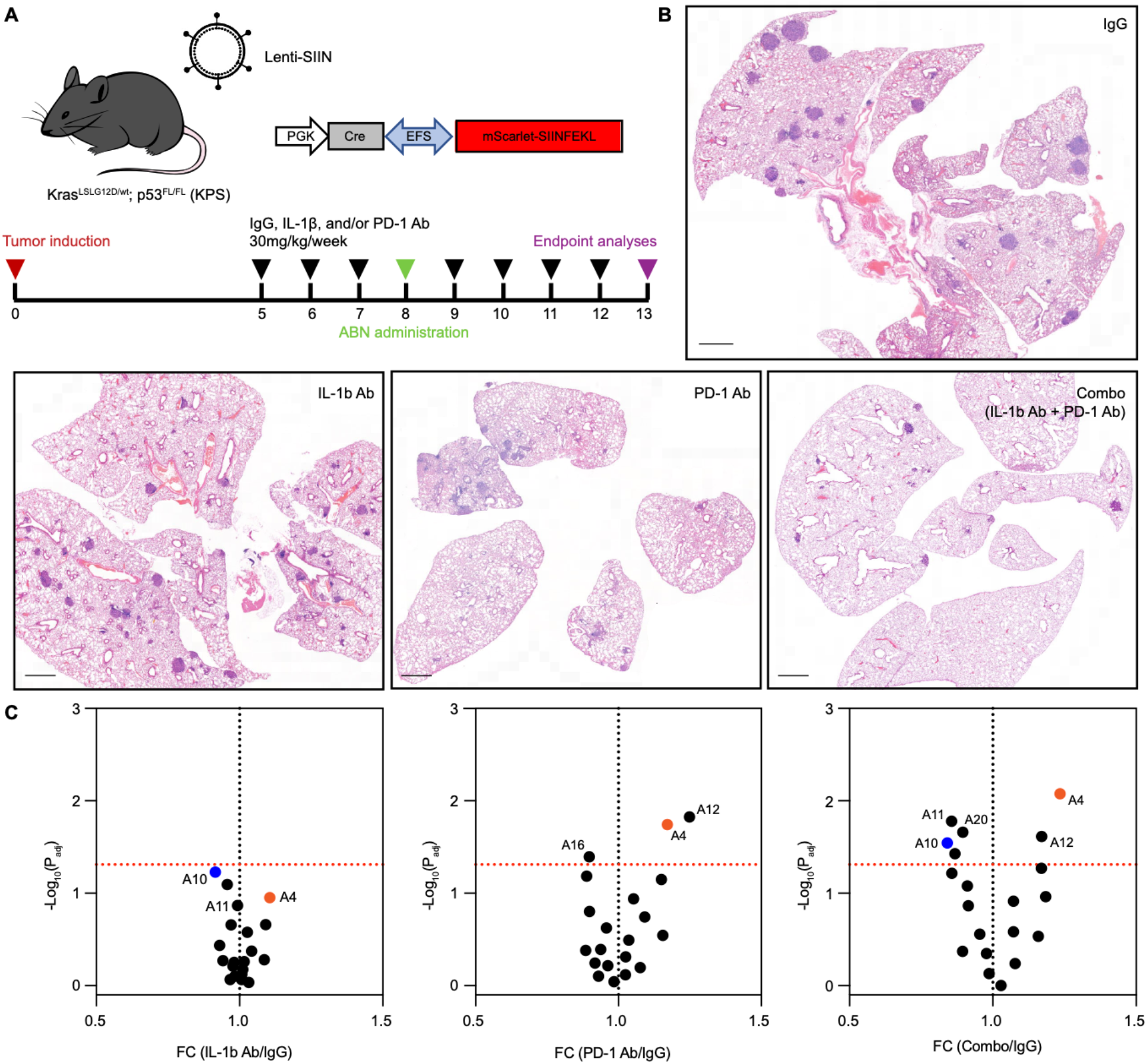
Protease activity differentiates treated versus untreated KPS lung adenocarcinoma. (**A**) KP mice were injected intratracheally with lentivirus encoding for Cre recombinase and the model antigen SIINFEKL (Lenti-SIIN, red triangle) to develop the inflamed “KPS” model. Weekly antibody treatments were started five weeks post tumor induction (black triangles). After three weeks of treatment, a panel of ABNs were administered and cleaved reporters were collected in urine (green triangle). After eight weeks of treatment, mice were sacrificed, and tissues were collected for endpoint analyses (purple triangle). (**B**) Histology of fixed lung tissue after treatment show tumor burdens under different therapeutic conditions (IgG control antibody, IL-1b antibody, PD-1 antibody, or combination of IL-1b and PD-1 antibodies). Scale bar 1mm. (**C**) Volcano plots of the fold changes of cleavage of the ABN panel after three weeks of treatment compared to IgG control. A4 (orange) and A10 (blue) substrates were found to be efficiently cleaved by granzyme B and caspase-1, respectively.

Using the same dosing scheme, we administered the 20-substrate panel of ABNs three weeks after initiating intervention treatment protocols (eight weeks after initiation of tumorigenesis) and collected urine to assess substrate cleavage. We found several differentially cleaved substrates at this time point (Fig. 2C). In particular, A10, sensitive to caspase-1 activity, exhibited the most significant reduction in cleavage fold change with IL-1b antibody treatment, relative to IgG control-treated tumor-bearing mice. With PD-1 antibody treatment, more sensor release was observed for the neutrophil elastase (ELANE) (A12) and granzyme B (A4) substrates, which is consistent with ongoing anti-tumor cytotoxic immunity in these treated mice. In the case of combination therapy, we saw both effects: increased ELANE and granzyme B activity in addition to decreased caspase-1 activity (A10 and A11).

### Immunofluorescence and AZP staining show IL-1b-associated proteases in KPS tissue

In addition to using ABNs to measure protease activity *in vivo*, we sought to evaluate the spatial abundance of IL-1b-associated proteases in the KPS mice. We compared immunofluorescence staining for caspase-1, cathepsin G (CTSG), and ELANE, which are known to be upstream mediators of IL-1b activation, in KPS lung tissue sections over time. We found high levels of caspase-1 and CTSG protein, as well as minimal expression of ELANE, in tumors that developed six weeks post-induction, and this pattern persisted through later stages (Fig. 3A, S3). As expression does not necessitate enzymatic activity, we turned to activity zymography probes (AZPs) as a tool to localize the observed protease cleavage activity within the sections (Fig. 3B). AZPs consist of a fluorophore-labeled positively charged domain linked to a masking agent via a protease-cleavable substrate sequence [18]. When this substrate is cleaved by local protease activity, the positively charged fluorescent fragment can bind to negatively charged tissue in the vicinity. The target substrates were chosen from our FRET screen and ABN panel, in which we had identified specifically and efficiently cleaved substrates for each protease of interest. A1, A10, and A12 are cleaved robustly by recombinant CTSG, caspase-1, and ELANE, respectively (Fig. 1C, S4, S5). AZP staining in KPS tumors showed high A10 signal but minimal A1 and A12 cleavage (Fig. 3C, Table S3). Co-incubation of AZPs with broad-spectrum inhibitors blocked enzymatic cleavage and showed no fluorescence signal (S6). These findings were consistent with the absence of ELANE expression observed in immunofluorescence and the lack of CTSG activity amongst the *in vivo* nanosensors. With further exploration of A10, we observed a robust fluorescence AZP signal that highlighted tumors in the KPS lung tissue. In contrast, significantly less A10 staining was detected in both normal adjacent tissue (NAT) and in healthy tissue samples, indicating tumor-specific cleavage and binding of the probe (Fig. 3D). Collectively, the detection trends of the A10 sensor were consistent with the hypothesis that caspase-1 activity is present during and potentially contributes to cancer progression in untreated KPS mice.

**Figure 3.**
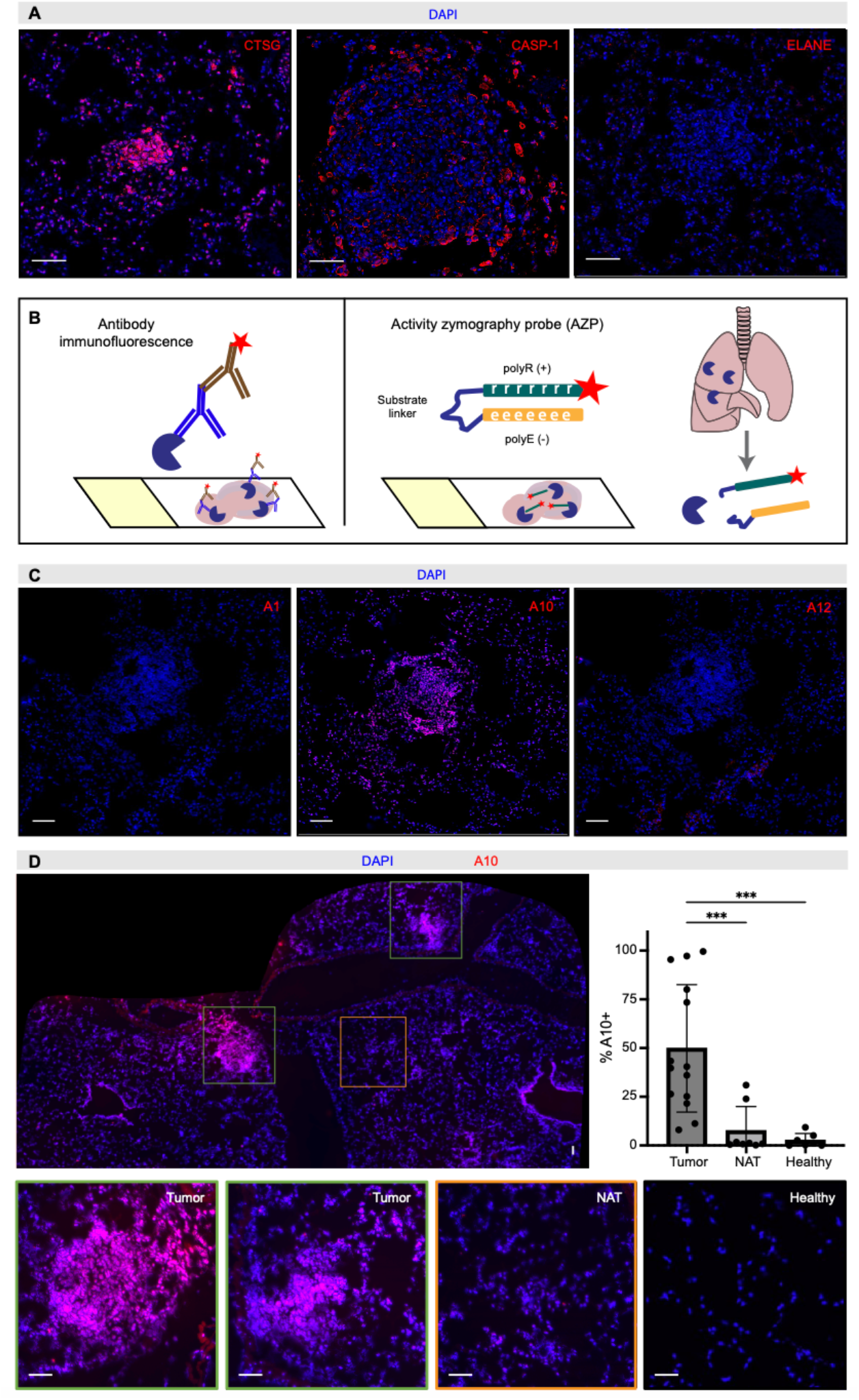
Caspase-1 is highly expressed and highlights tumors in an activity-dependent manner. (**A**) Immunofluorescence staining of protease protein expression (red): cathepsin G (CTSG), caspase-1, and neutrophil elastase (ELANE), in 6-week KPS tumors. All sections counterstained with DAPI (blue). Scale bar 50um. (**B**) Schematic of antibody versus activity zymography probe (AZP) staining on frozen tissue sections. Protease cleavable substrates can be converted to AZPs that show cleavage locally on tissue. AZPs consist of a negatively charged reporter masked by a peptide substrate sequence that prevents it from binding to tissue until cleaved by localized target proteases. (**C**) Binding of activated AZPs (red) cleavable by CTSG (A1), caspase-1 (A10), and ELANE (A12) in lung tumor tissue. All sections counterstained with DAPI (blue). Scale bar 50um. (**D**) A10 AZP stained 7-week KPS lung sections significantly more in tumors and less in normal adjacent tissue (NAT) or healthy tissue. All sections counterstained with DAPI (blue). Scale bar 25um. Percentage of tissue regions that were positively stained for A10 were quantified. One-way ANOVA with Tukey’s multiple comparison test was performed (p < 0.05).

### A10 highlights caspase-1 activity in lung tumors

Co-staining tumor sections with the A10 AZP and a caspase-1 antibody showed protein expression co-localizes with sensor cleavage in both early- and late-stage KPS tumors (Fig. 4A, S7). The A10 activity-based probe’s staining pattern does not exactly overlap with the antibody’s immunofluorescent binding, likely due to the limited resolution of probe binding. In this case, upon activation by enzymatic cleavage, the AZP’s positive probe fraction is liberated and seeks to bind to nearby, negatively charged nuclei as the strongest electrostatic interaction. This outcome results in regional localization, but a lower cellular staining resolution compared to antibodies, for which the fluorescent label binds directly to a specific protein epitope. When the tissue samples exposed to the A10 AZP were co-incubated with a broad-spectrum protease inhibitor or a metabolized caspase-1-specific inhibitor, VRT-043198, we observed a significant reduction in fluorescence labeling of the tumors (Fig. 4B). This inhibition-dependent decrease in tumor staining suggested that caspase-1 activity was significantly contributing to local tissue cleavage of A10. We further validated the role of caspase-1 in cleaving the A10 substrate using lung homogenates from late-stage KPS mice in a FRET inhibitor assay (Fig. 4C, S8). We found that most of the cleavage signal produced by the tumor-bearing lung tissue samples was due to caspase-1, given that the fold change in fluorescence signal was significantly lower when tumor homogenate samples were incubated with the caspase-1-specific inhibitor.

**Figure 4.**
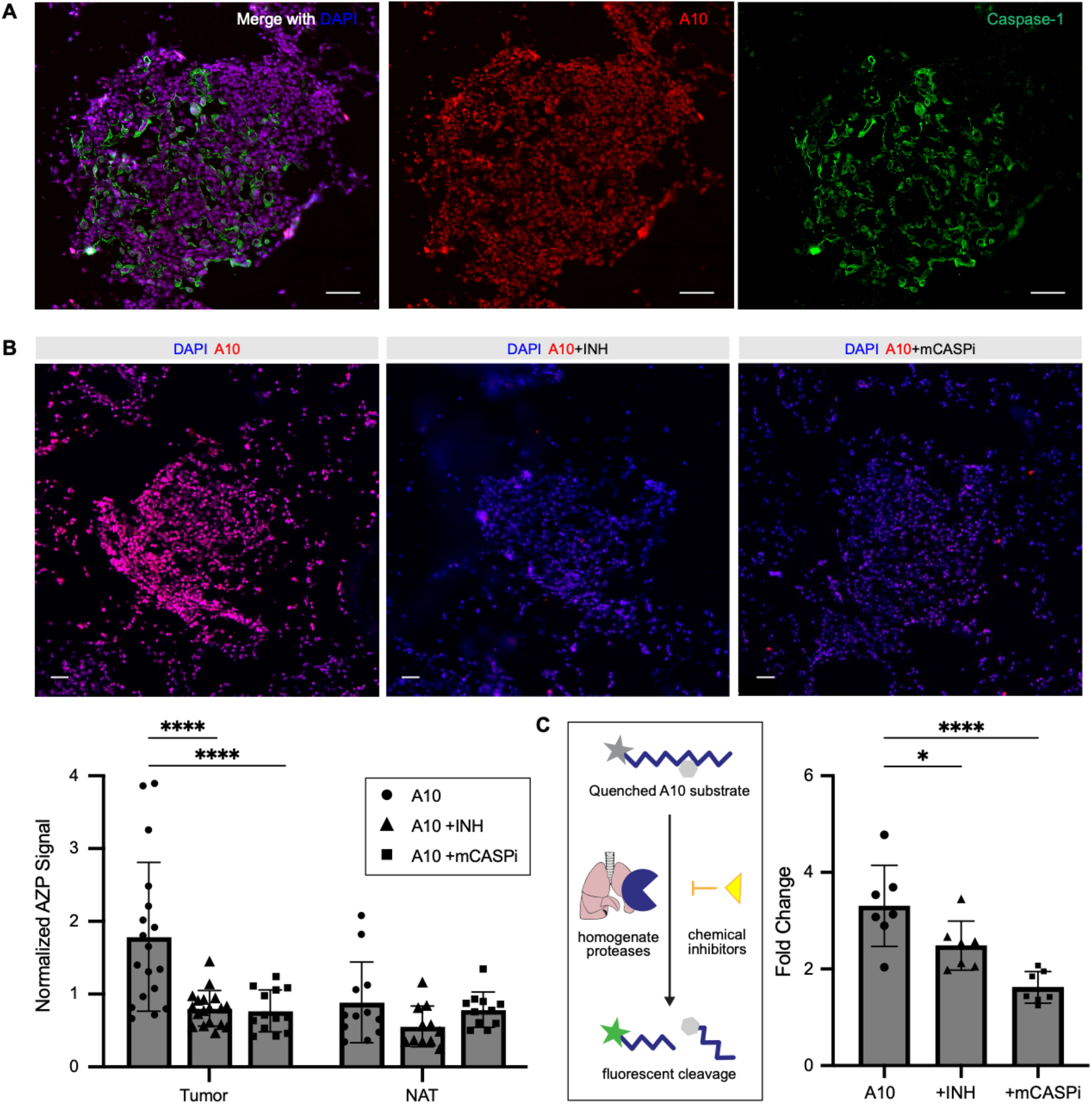
A10 AZP cleavage is abrogated by caspase-1 inhibition. (**A**) Co-staining of caspase-1 with A10 AZP in 7-week KPS tumor. Scale bar 50um. (**B**) A10 AZP stained 7-week KPS lung sections were incubated with broad-spectrum (INH) or metabolized caspase-1-specific inhibitor (mCASPi). All sections counterstained with DAPI (blue). Normalized A10 AZP signals in each tumor region of a given mouse were quantified and compared using one-way ANOVA with Tukey’s multiple comparison test was performed (p < 0.05). Scale bar 25um. (**C**) 11-week KPS lung homogenates were incubated with A10 FRET probe, INH, and/or mCASPi, where fold changes indicate increase in fluorescence from cleavage after 120 min. One-way ANOVA with Tukey’s multiple comparison test was performed (p < 0.05).

### Human lung adenocarcinoma samples exhibit high caspase-1 expression and activity

Beyond the KPS model, we explored the translational relevance of caspase-1 by examining human LUAD samples. Immunohistochemical staining of caspase-1 in sections of human lung biopsies showed elevated numbers of caspase-1-positive cells in tumor areas, relative to adjacent healthy areas (S9). We also tested bronchioalveolar lavage fluid (BALF), which is a common form of clinical samples that capture interstitial cells and proteins from the lungs. We hypothesized that caspase-1 would be released and active in this compartment in samples from LUAD patients. BALF samples were procured from healthy smokers with no detectable cancer, as well as from smokers who were patients with Stage I/II LUAD. The BALF samples, normalized by total protein concentration, were assessed for caspase-1 content via ELISA. There was no detectable caspase-1 in healthy BALF, in contrast to varying levels of caspase-1 in cancer samples (S10a). To evaluate caspase-1 activity, we incubated the BALF samples with quenched luciferase-conjugated caspase-1 substrates, where high luminescence indicated that active caspase-1 from the samples had recognized and cleaved the substrate. BALF samples from patients with LUAD had significantly higher luminescent signals (S10b). This observation implies that caspase-1 expression and activity may be a result of cancer-specific processes, rather than due to inflammation from smoking.

### Combined blockade of IL-1b and caspase-1 significantly reduces lung tumor formation

As caspase-1 expression and activity have been shown to be elevated in both murine and human tumor samples, we hypothesized that caspase-1 may act as an upstream activator of pro-tumor inflammation and that we could therapeutically perturb it to reduce tumorigenesis. We applied a small molecule caspase-1 inhibitor, belnacasan, at a daily oral gavage dose of 100 mg/kg [25]. We combined belnacasan administration with the IL-1b antibody as a novel treatment strategy and compared it with control IgG antibody or IL-1b antibody alone (Fig. 5A). We started administering the candidate therapeutics to KPS mice four weeks after tumor inoculation. After seven weeks of treatment, we detected significantly fewer number of tumor nodules when belnacasan was administered, and this improvement was even greater when combined with IL-1b antibody blockade as shown by both lung cross-section histology and volumetric microCT quantification (Fig. 5B-D, S11). Moreover, the total tumor burden was also reduced in the caspase-1-inhibited cohort, and the individual tumor nodules that were detected in treated mice were significantly smaller. Notably, 18% of combination-treated KPS mice bore no visible tumors, highlighting a dramatic example of lung tumor interception. Immunohistochemical staining of active IL-1b in sections of treated lungs further shows decreased IL-1b and local reduction of inflammation (S12). While IL-1b antibody alone lacked generalizable therapeutic efficacy, as seen by the wide distribution spread in tumor burden, the combined blockade with caspase-1 inhibition was able to more comprehensively block both the activation and pro-inflammatory function of IL-1b, leading to a drastically improved response rate (Fig. 5E).

**Figure 5.**
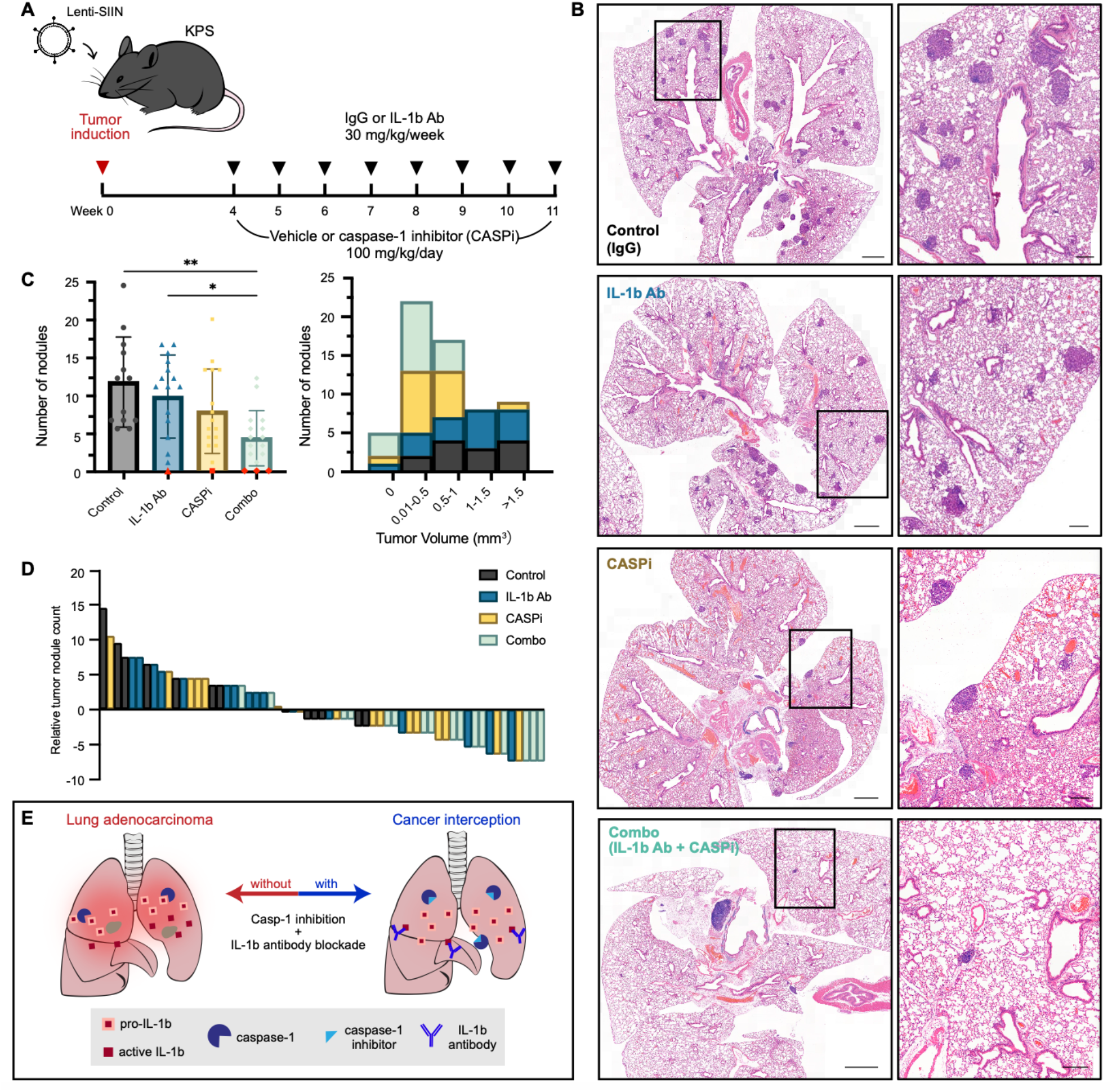
Combined caspase-1 inhibition and IL-1b blockade reduces KPS lung tumor burden. (**A**) KPS mice were induced with lentivirus encoding for Cre recombinase and the model antigen SIINFEKL. Weekly antibody treatments (30mg/kg) and daily oral gavage of the caspase-1 inhibitor belnacasan (CASPi, 100mg/kg) were started four weeks post-tumor induction. (**B**) Histology for representative lungs from in each treatment group (scale bar 1mm) with enlarged insets (scale bar 250um). (**C**) Tumor burden volume and (**D**) number of nodules were quantified from microCT images (n=13-16 from two independent cohorts). Mice with no detectable nodules are marked with red circles (**C, left**) or listed as having 0 tumor volume. One-way ANOVA with Tukey’s multiple comparison test was performed (p < 0.05). (**E**) Schematic of lung cancer interception leveraging combined caspase-1 inhibition and IL-1b blockade to reduce inflammation.

## Discussion

Here, we integrated activity-based nanosensors across hierarchical scales to probe IL-1b-associated proteolytic activity and its contribution to the progression of inflammatory LUAD. We uncovered caspase-1 as a highly expressed and active upstream regulator of IL-1b in KPS tumors and developed a novel therapeutic approach that combined IL-1b blockade with caspase-1 inhibition to improve cancer outcomes.

IL-1b antibody (canakinumab) treatment was reported to reduce the overall cancer mortality and incidence of lung cancer in patients in the CANTOS trial, but there was little improvement in survival when a similar therapeutic regimen was applied for advanced or metastatic NSCLC [2]. These observations suggest that knowledge gaps surrounding IL-1b activation and its role in the TME are hindering the development of a more effective therapeutic approach in LUAD. We were able to use a representative inflammatory LUAD murine model to make advances in understanding IL-1b activation and its role in the TME. Leveraging potential field cancerization in the lungs, our ABN platform is able to capture and monitor dysregulated protease activity in the local lung environment *in vivo* [16, 23, 26]. We designed a panel using substrates that were previously identified to differentiate lung cancer from healthy mice and new substrates that we screened against upstream activators of IL-1b [14, 17]. We observed increased cleavage of a substrate susceptible to granzyme B activity upon treatment, consistent with its role as a mediator of cytotoxic lymphocyte activity [27]. In contrast, among all the proteases we probed that are involved in IL-1b activation, caspase-1 activity decreased most significantly with treatment and tumor cell death, thus appearing to contribute the most to the cancer state. Caspase-1 exerts a dichotomous influence in cancer through the inflammasome pathway: pyroptosis can kill tumor cells, but IL-1b activation can support tumor growth and immune escape during chronic inflammation [28, 29].

In addition to LUAD, inflammation from IL-1b signaling is also implicated in hematological malignancies and cardiovascular diseases (CVD). Enriched levels of inflammasome components that activate caspase-1, as well as IL-1b, were found in macrophages in atherosclerotic CVD [30, 31]. Accumulation of IL-1b+ macrophages in tumors have been found to contribute to TME remodeling, metastasis, and therapeutic resistance [32]. While we did not pinpoint the specific cell source for caspase-1 production and activation in LUAD progression, there is generally a high tumor infiltration of immune cells in KPS tumors, including neutrophils, T cells, and macrophages [13]. Tumor-infiltrating clonal hematopoiesis has also been found to increase the risk of NSCLC-associated death by remodeling the tumor immune microenvironment, highlighting patients with these mutations as a potential population that could benefit from our findings [33]. Cellular analysis, such as with single cell RNA sequencing, for LUAD might be able to elucidate the mechanistic caspase-1-associated genetic programs in these different cell populations that result in tumor development. The balance of immunosuppressive and immune-activating populations would also merit further exploration.

To more robustly block the pro-tumorigenic IL-1b pathway via the upstream caspase-1, we proposed a novel therapeutic strategy combining IL-1b antibody and caspase-1 inhibitor, belnacasan. Belnacasan, or VX-765, is an oral pro-drug found to reduce immune activation and delay pathology in inflammatory diseases [34]. It has been used in clinical trials for psoriasis, epilepsy, and COVID-19 (NCT01048255, NCT00205465, NCT01501383, NCT05164120). Although it was well-tolerated and safe for administration in humans, it lacked therapeutic efficacy in these prior indications. Cancer applications of belnacasan are still being explored; one recent example includes its use in mice to improve sensitivity to immune checkpoint blockade in triple-negative breast cancer xenografts [35]. In this study, we demonstrated that belnacasan could be repurposed for lung cancer interception. When applied at an early stage, caspase-1 inhibition alone significantly reduced tumor burden in KPS mice. Outcomes were reliably and further improved with the combination of IL-1b blockade, as indicated by the lower count and smaller size distribution of detected tumor nodules. This therapeutic impact was achieved potentially through both reducing the activation of IL-1b and blocking the downstream functions of any active IL-1b. While we were able to identify caspase-1 activity as an early carcinogenic event and therapeutic target in LUAD, future work to understand how other inflammatory factors change over time with progression would help determine the stage of intervention and patient stratification strategies to bring this finding closer to human application [36]. Furthermore, investigating the antigen-specific immune response from this combination treatment will be enlightening to understand resulting adaptive immunity and long-term memory. Given inflammation’s far-reaching role in various other diseases, such as ulcerative colitis and rheumatoid arthritis, combination therapies that combine caspase-1 inhibition and a more disease-specific target may also be promising to explore more broadly.

Our study investigated and monitored IL-1b-associated protease dysfunction to seek mechanistic insights into inflammatory lung cancer. The spatial and longitudinal characterization of proteases in the tumor microenvironment through various nanoscale modalities shed light on pro-tumorigenic features of the lung TME as potential translational targets for cancer interception, amidst the crosstalk of inflammation and tumorigenesis in LUAD. Combination therapy of IL-1b blockade with caspase-1 inhibition showed significant reduction in tumor burden, suggesting a potential path to cancer interception at-risk patients with prolonged exposure to smoke or other air pollutants. We envision that early administration of this therapy in high-risk patient populations could transform the lung cancer interception landscape. More broadly, our activity-based nanotechnologies serve as a widely applicable way to probe and better discover disease biology.

## Materials and Methods

### Fluorescence Resonance Energy Transfer (FRET) assay

FRET probes were ordered from CPC Scientific (Sunnyvale, CA) and designed to have the quencher 2,4-dinitrophenyl-lysine (DNP) and fluorophore 7-Methoxycoumarin-4-acetic acid (MCA) or 6-carboxyfluorescein (FAM) on either side of the cleavable peptide sequences. FRET probes (50 uM) were mixed with recombinant proteins (10 nM) or lung homogenate (total protein concentration: 1 mg/ml) to initiate the cleavage reaction, and the fluorescence signal was read out periodically with a Tecan plate reader for over an hour. Recombinant proteases screened included caspase-1 (ALX-201-056-U025, Enzo), caspase-3 (707-C3, R&D Systems), caspase-7 (823-C7, R&D Systems), caspase-8 (ALX-201-163-C020, Enzo), cathepsin B (965-CY, R&D Systems), cathepsin G (BML-SE283-0100, Enzo), chymase-1 (4099-SE, R&D Systems), chymotrypsin (6907-SE, R&D Systems), granzyme B (1865-SE, R&D Systems), meprin A (4007-ZN, R&D Systems), meprin B (3300-ZN, R&D Systems), and neutrophil elastase (4517-SE, R&D Systems). For FRET with lung homogenate samples, chemical inhibitors were added to validate protease activity-enabled fluorescence. 100x dilution of Halt™ Protease Inhibitor Cocktail (78429, ThermoFisher Scientific) and 1 mM marimastat (HY-12169, MedChemExpress) were used together for pan-protease inhibition. 1 mM VRT-043198 (HY-112226, MedChemExpress) was used for caspase-1-specific inhibition.

### KPS murine model

To develop the KPS LUAD murine model, we generated KrasLSL-G12D;Trp53fl/fl C57BL/6 (KP) mice in which the activation of an oncogenic allele of Kras is sufficient to initiate the tumorigenesis process, and additional deletion or point mutation of p53 substantially enhances tumor progression [37]. LUAD lung tumors were initiated in KP mice that were 8-12 weeks old by intratracheal administration of 10,000 plaque-forming units of lentivirus expressing Cre-recombinase and a tumor-specific neoantigen SIINFEKL fused to the fluorophore mScarlet (Lenti-SIIN) [13]. The mice receiving Lenti-SIIN were then randomly grouped to minimize the variation in viral delivery. Healthy control cohorts consisted of age- and sex-matched mice. Tumor growth was allowed in KPS mice until the treatment or detection was performed. During selection of KPS mice for treatment groups, investigators were blinded to all characteristics but age, sex, and genotype. All animal studies and procedures were approved by the Committee for Animal Care at Massachusetts Institute of Technology (MIT) (protocols 2301000462, 2212000452) and were conducted in compliance with institutional and national policies. All animal studies were performed in compliance with Animal Research: Reporting In Vivo Experiments (ARRIVE) guidelines.

### *In vivo* treatment studies

To evaluate IL-1b blockade in KPS mice, all tumor-bearing mice received one of the following treatment regimens at the end of five weeks post LUAD initiation: IgG control, anti-IL-1b, anti-PD-1, and anti-IL-1b with anti-PD-1. The antibodies, anti-Armenian hamster IgG (BE0091, InVivoMAb), anti-mouse/rat IL-1b (BE0246, InVivoMAb), and anti-mouse PD-1 (BP0273, InVivoMAb), were administered three times a week intraperitoneally at 10 mg/kg. The treatment continued for nine weeks, with longitudinal monitoring of tumor growth by microCT every two or three weeks and ABN administration after three weeks of treatment.

To test caspase-1 inhibition *in vivo*, KPS mice were treated four weeks post lentiviral induction. The treatment groups were IgG with drug vehicle, anti-IL-1b with drug vehicle, IgG with belnacasan, and anti-IL-1b with belnacasan. Mice were dosed with antibody intraperitoneally three times a week at 10 mg/kg. Belnacasan (HY-13205, MedChemExpress) was solubilized in the vehicle (20% Kolliphor in water) at a concentration of 200 mg/ml. Mice were administered 100 mg/kg belnacasan or vehicle control every day via oral gavage, starting on the same day as the antibody treatment. The treatment continued for seven weeks, with longitudinal monitoring of tumor growth by microCT every two or three weeks.

### Intratracheal ABN administration

Mass-encoded ABNs were synthesized and characterized by CPC Scientific (Sunnyvale, CA), consisting of glutamate-fibrinopeptide B (GluFib, EGVNDNEEGFFSAR) reporters with different distributions of stable isotopes linked to an 8-arm 40kDa PEG via unique amino acid sequence linkers. For intratracheal instillation, a 20-plexed mass-encoded ABN cocktail (20 uM substrate) in phosphate buffer (0.28 M mannitol, 5 mM sodium phosphate monobasic, 15 mM sodium phosphate dibasic, pH 7.2 to 7.4) was administered by solution injection following intratracheal intubation with a 22-gauge flexible plastic catheter (Exel International, Redondo Beach, CA). 100 ul of air was then pushed into mouse lungs via a 1-ml syringe to ensure a full delivery of ABNs into the lungs. A subcutaneous injection of 200 ul sterile PBS was administered in favor of urine production. Bladders were voided at 60 min after nanosensor administration, and urines produced from 60 to 120 min after administration were collected. All urine samples were immediately frozen at −80°C until analysis by a quadruple LC-MS/MS in the Biopolymers and Proteomic Core at the Koch Institute for Integrative Cancer Research.

### Lung tissue collection and preparation

For frozen tissue sections, mouse lungs were collected and embedded in OCT blocks. OCT blocks were frozen over dry ice and stored at -80C for cryosectioning. For histology, collected mouse lungs were fixed in 4% paraformaldehyde before paraffin-embedding for sectioning and H&E staining. BALF samples were collected by intratracheally instilling 1 ml of PBS and suctioning the contents. They were then spun down and removed of cell pellet.

### Immunofluorescence and activatable zymography probe (AZP) staining of frozen tissue

The cleavable peptide sequence was ordered from CPC Scientific, flanked by a negatively charged E9 domain and a positively charged R9 domain, with the Cy5 fluorophore conjugated to the end of the positive domain. A polyR peptide consisting of the R9 domain with a Cy7 fluorophore was used as a positive control for electrostatic binding. Staining with AZPs was performed as previously described [18]. Briefly, frozen tissue sections were prepared by washing with PBS and fixing with ice-cold acetone. They were then blocked with 1% BSA at room temperature before incubating with the AZP probes and primary antibodies, if applicable, for 4 hours at 37°C. For studies with only antibody staining, tissues were blocked in the same manner and incubated with antibodies overnight at 4°C. The tissues were then washed with PBS and incubated with secondary antibodies diluted in PBS for 30 minutes at room temperature. Primary antibodies used include caspase-1 (22915-1-AP, ThermoFisher Scientific), cathepsin G (PA5-99402, ThermoFisher Scientific), and neutrophil elastase (PA5-115648, ThermoFisher Scientific). For studies including tissue protease inhibition, inhibitors were co-incubated with the blocking solution and with the AZP probes. 100x dilution of Halt™ Protease Inhibitor Cocktail (78429, ThermoFisher Scientific), 1 mM marimastat (HY-12169, MedChemExpress), and 1 mM 4-(2-Aminoethyl)benzenesulfonyl fluoride hydrochloride (AEBSF) (A8456, Millipore Sigma) were used together for pan-protease inhibition. 1 mM belnacasan (HY-13205, MedChemExpress) was used for caspase-1-specific inhibition. For quantification, cells with fluorescence intensity signals above a pre-determined threshold were detected as antibody marker+ using QuPath 0.5.1 [38]. AZP signals were quantified by normalizing the AZP Cy5 signal with the polyR Cy7 signal for each cell.

### Immunohistochemical staining of fixed tissue

Slides with FFPE tissue sections were deparaffinized with xylene, rehydrated, and antigen-retrieved in at 97°C for 20 minutes using HIER citrate buffer (pH 6). They were then blocked for 30 minutes, followed by incubation with primary antibody for one hour, horseradish peroxidase (HRP) for 30 minutes, and DAB substrate for five minutes. Primary antibodies for IL-1b (218753, Bio-X-Cell) and caspase-1 (MA5-16215, ThermoFisher Scientific) were diluted with Tris buffered saline with Tween-20, and HRP was used for secondary detection.

### Human LUAD sample processing

Sections from fresh frozen human LUAD samples and normal adjacent tissue were acquired from the Massachusetts General Hospital Lung Tumor Bank. De-identified BALF samples from patients with LUAD and from healthy donors were purchased from Bay Biosciences LLC (Brookline, MA). Human BALF samples were assessed for human caspase-1 expression with a solid phase sandwich enzyme-linked immunosorbent assay (ELISA) kit (DCA100, R&D Systems), as described via kit protocol. Caspase-1 activity in BALF was assessed with a bioluminescence-based assay, as described via kit protocol (G9951, Promega). Cells with positive staining for DAB were detected and quantified with QuPath 0.5.1 [38].

### Statistics

Statistical analyses were performed using GraphPad Prism v10. One-way ANOVA with Tukey’s multiple comparison test was used for statistical testing unless noted otherwise. Statistical significance is indicated as *P < 0.05, **P < 0.01, ***P < 0.005, and ****P < 0.001.

## Acknowledgments

We thank Koch Institute Swanson Biotechnology Center, specifically the Histology core; Dr. I. Dagogo-Jack (Massachusetts General Hospital, Boston, MA) and the MGH Lung Tumor Bank for the de-identified human tissues. This study was supported, in part, by Johnson & Johnson, the Virginia and D.K. Ludwig Fund for Cancer Research, the Koch Institute Frontier Research Program through a gift from Upstage Lung Cancer, and the Koch Institute’s Marble Center for Cancer Nanomedicine. C.S.W. acknowledges support from the National Science Foundation Graduate Research Fellowship and the Graduate Student Fellowship from the Ludwig Center at MIT‘s Koch Institute. C.M.A. acknowledges support from La Caixa Foundation and the Graduate Student Fellowship from the Ludwig Center at MIT‘s Koch Institute. S.P. acknowledges support from National Institute of Health T32 (T32HL116275). S.N.B. is a Howard Hughes Medical Institute Investigator.

## Competing interests

A provisional patent has been filed on this work: “Profiling inflammation-associated protease activity in early lung cancer” (CSW, SNB). SNB reports consulting roles, board membership, and/or equity in Global Oncology, Sunbird Bio, Satellite Bio, Matrisome Bio, Xilio Therapeutics, Danaher, Catalio Capital, Pictet, Ochre Bio, Amplifyer Bio, Earli Inc., Impilo Therapeutics, Port Therapeutics, Vertex Pharmaceuticals, Ropirio Therapeutics, and Moderna and receives sponsored research funding from Open Philanthropy and Owlstone Medical, which were not involved in this study. TEJ reports consulting roles, board membership, and/or equity in Amgen, Thermo Fisher Scientific, Dragonfly Therapeutics, T2 Biosystems, SQZ Biotech, Skyhawk Therapeutics, and Break Through Cancer and receives sponsored research funding from the Lustgarten Foundation, which were not involved in this study. The remaining authors report no competing interests.

## Data and materials availability

All data needed to evaluate the conclusions in the paper are present in the paper and/or the Supplementary Materials.

