## Supplementary Information for "Multimodal profiling of proinflammatory protease activity identifies caspase-1 as a target for lung cancer interception"

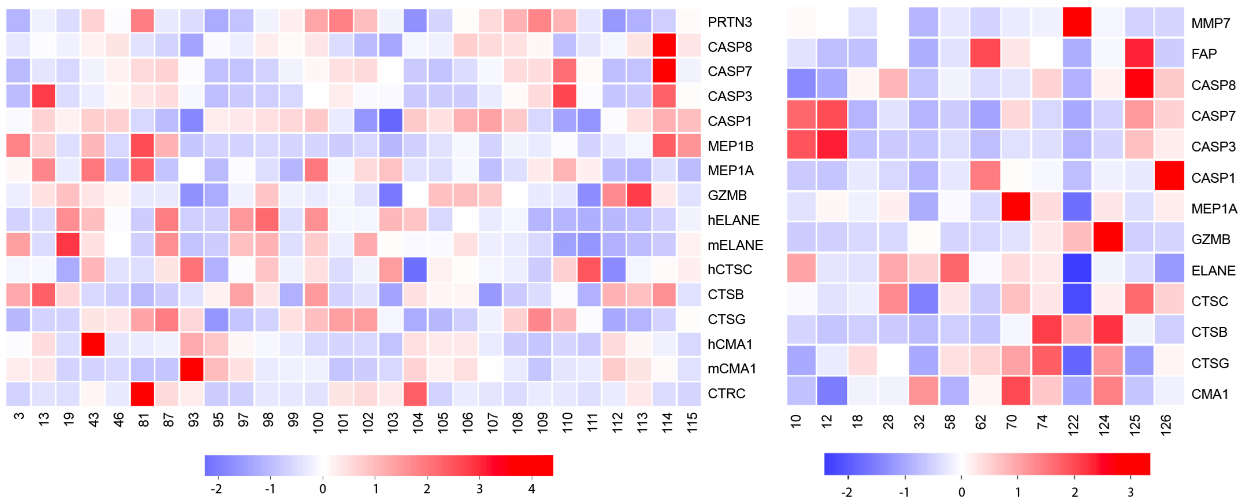


**Figure S1. Peptide substrate cleavage screening.** Heatmaps depicting z-scored cleavage fold change of FRET substrates (columns) by interleukin- or cancer-associated proteases (rows). Proteases are mouse recombinant proteins unless noted otherwise. PRTN3: proteinase-3, CASP1/3/7/8: caspase-1/3/7/8, MEP1A/B: meprin A subunit alpha/beta; GZMB: granzyme B, h/mELANE: human/mouse neutrophil elastase, CTSB/C/G: cathepsin B/C/G; h/mCMA1: human/mouse chymase, CTRC: chymotrypsin C, MMP7: matrix metalloprotease-7, FAP: fibroblast activation protein

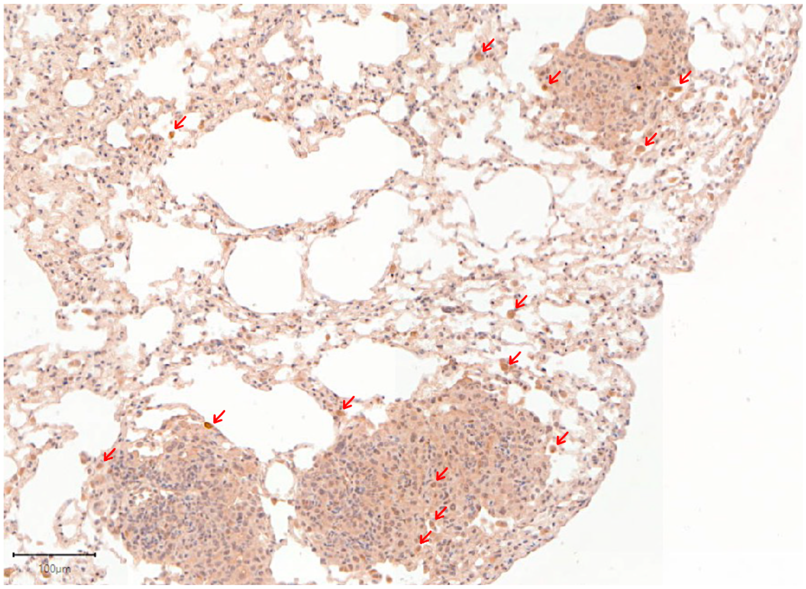


**Figure S2. Immunohistochemical staining of IL-1b.** Dark brown stained cells (examples shown by red arrows) show IL-1b expression in 12-week KPS lung tissue. Scale bar 100um.

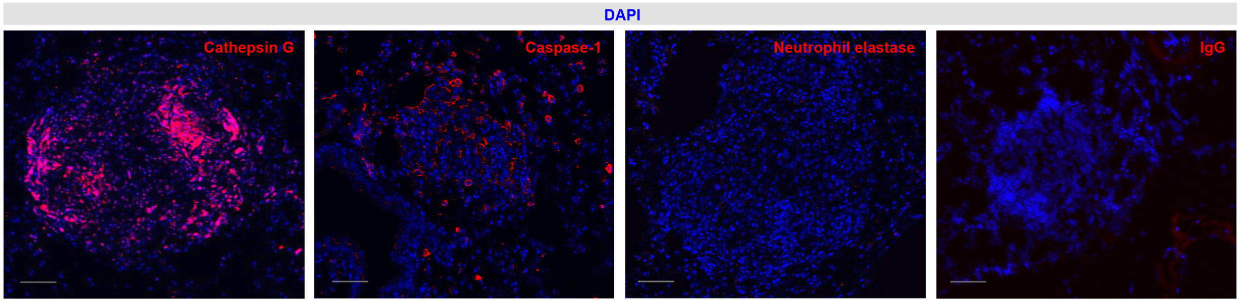


**Figure S3. Immunofluorescence staining of proteases.** Cathepsin G, caspase-1, neutrophil elastase, and IgG (red) are each co-stained with a nuclear dye (DAPI, blue) in 12-week KPS tumors. Scale bar 50um.

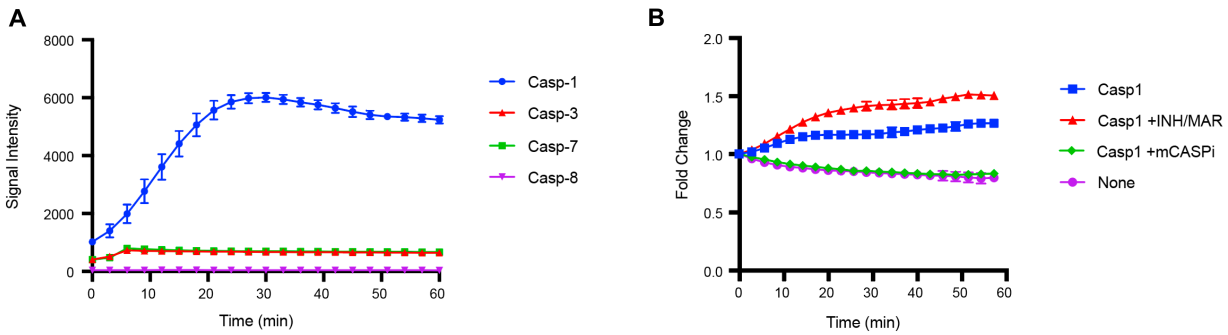


**Figure S4. Caspase-specific cleavage of A10 FRET probe.** (**A**) A10 was incubated with recombinant caspase-1 (blue), -3 (red), -7 (green), -8 (purple), where fluorescence would indicate FRET probe cleavage. N = 3 replicates. (**B**) A10 was incubated for cleavage by recombinant caspase-1 (blue), caspase-1 with broad spectrum inhibitor (INH) and marimastat (MAR) (red), or caspase-1 with metabolized caspase-1 inhibitor (mCASPi, green). A10 alone (None, purple) was used as a negative control. N = 3 replicates.

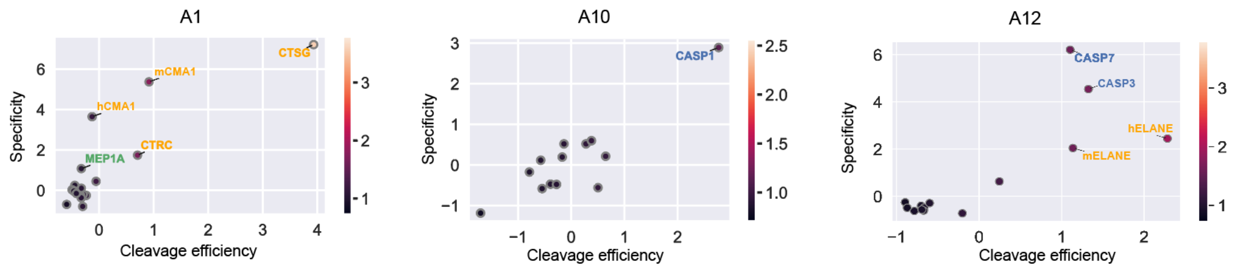


**Figure S5. Cleavage efficiency versus specificity plots for substrates of interest.** Standardized cleavage fold changes of different proteases are compared for A1, A10, and A12. A1 is highly cleaved by cathepsin G (CTSG), A10 by caspase-1 (CASP1), and A12 by neutrophil elastase (ELANE), and caspase-3/7 (CASP3/7). h-: human recombinant protein, m-: mouse recombinant protein. Text color indicates protease class, where green: matrix metalloprotease, yellow: serine, blue: cytosine.

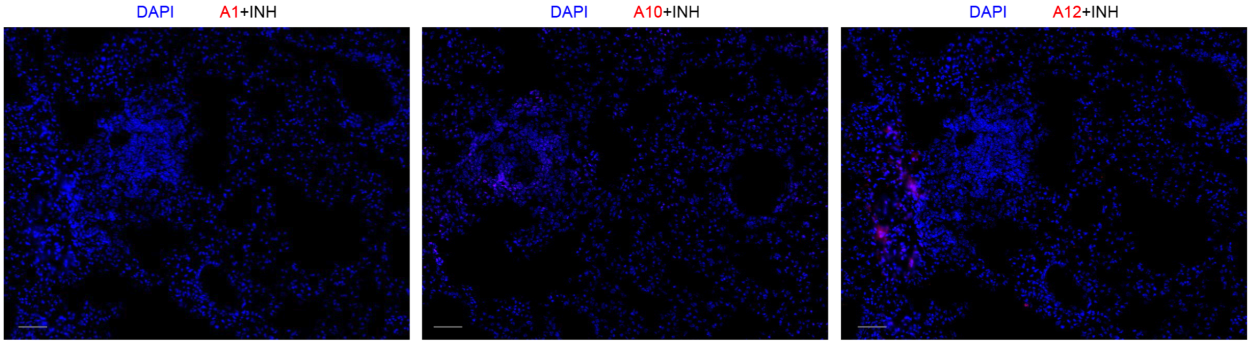


**Figure S6. Co-staining of AZPs with inhibitors on KPS tissue.** A1, A10, and A12 AZPs (red) are each incubated with broad spectrum protease inhibitor (INH) on 6-week KPS tissue to show cleavage-specific staining. Scale bar 50um.

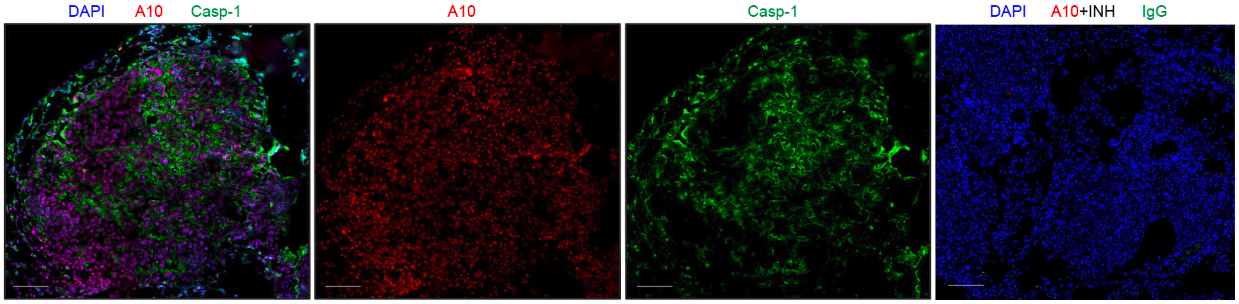


**Figure S7. Caspase-1 protein expression and activity staining.** Overlayed and single channel images of nuclear dye (DAPI, blue), AZP (A10, red), and caspase-1 (green) in 12-week KPS tumors. Scale bar 50um.

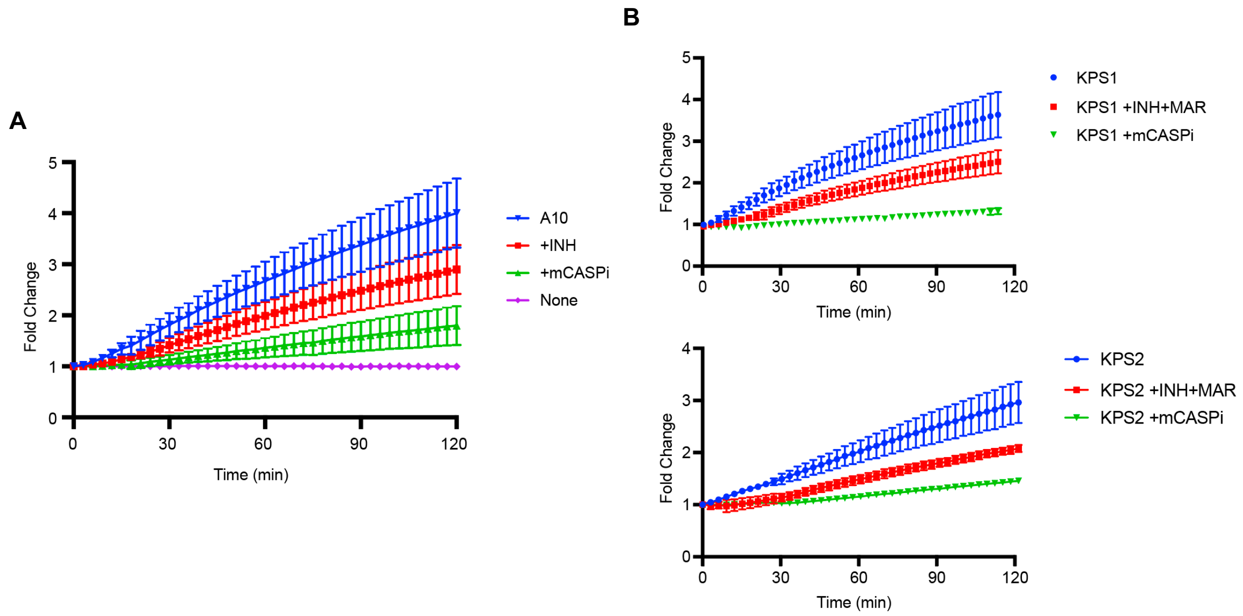


**Figure S8. FRET cleavage of A10 with lung homogenates from 12-week-old KPS mice.** (**A**) Fold change in cleavage of A10 FRET probe with and without broad spectrum inhibitor (INH) and marimastat (MAR) or metabolized caspase-1 inhibitor (mCASPi). N = 3 mice. (**B**) Examples of probe cleavage in individual mice. N = 2-3 replicates.

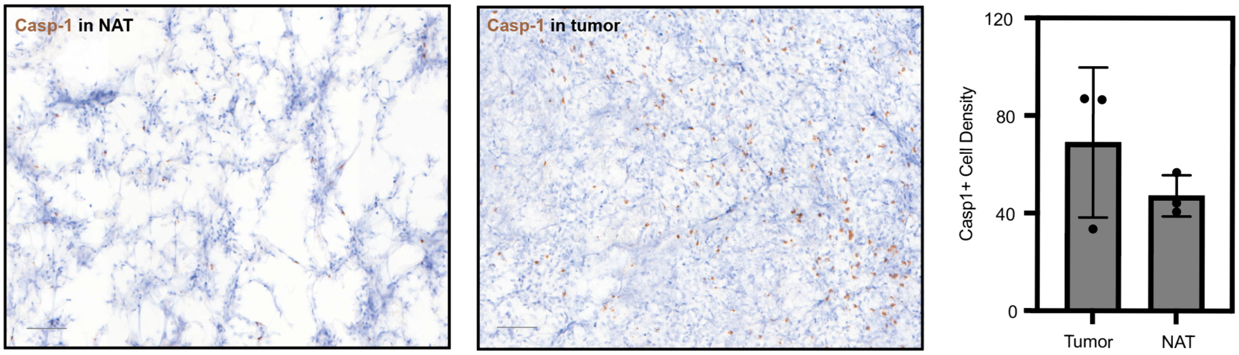


**Figure S9. Immunohistochemical staining of caspase-1 in human LUAD biopsy samples.** Caspase-1 (brown) expression was compared between normal tissue adjacent to tumor (NAT) and tumor sections. Scale bar 100um. Positive cell density for caspase-1+ expression was quantified in three separate sets of donor samples. N = 3 donor samples.

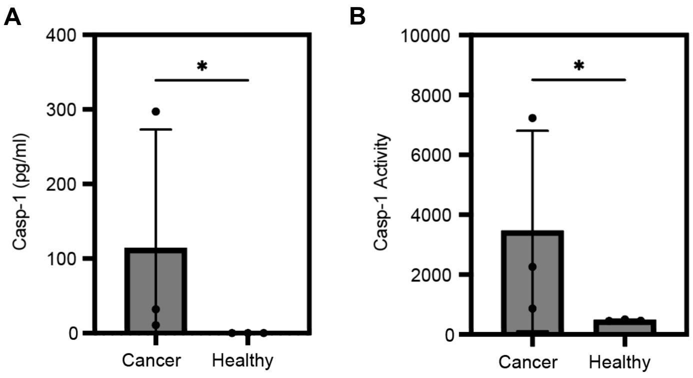


**Figure S10. Characterization of caspase-1 expression and activity in human bronchioalveolar lavage fluid (BALF).** (A) Caspase-1 expression in BALF from LUAD patients and from healthy smokers was quantified by ELISA. Mann-Whitney U test was performed ($p = 0.05$). N = 3 donor samples. (B) Caspase-1 activity in BALF was assessed with a commercial luminescence kit. Mann-Whitney U test was performed ($p = 0.05$). N = 3 donor samples.

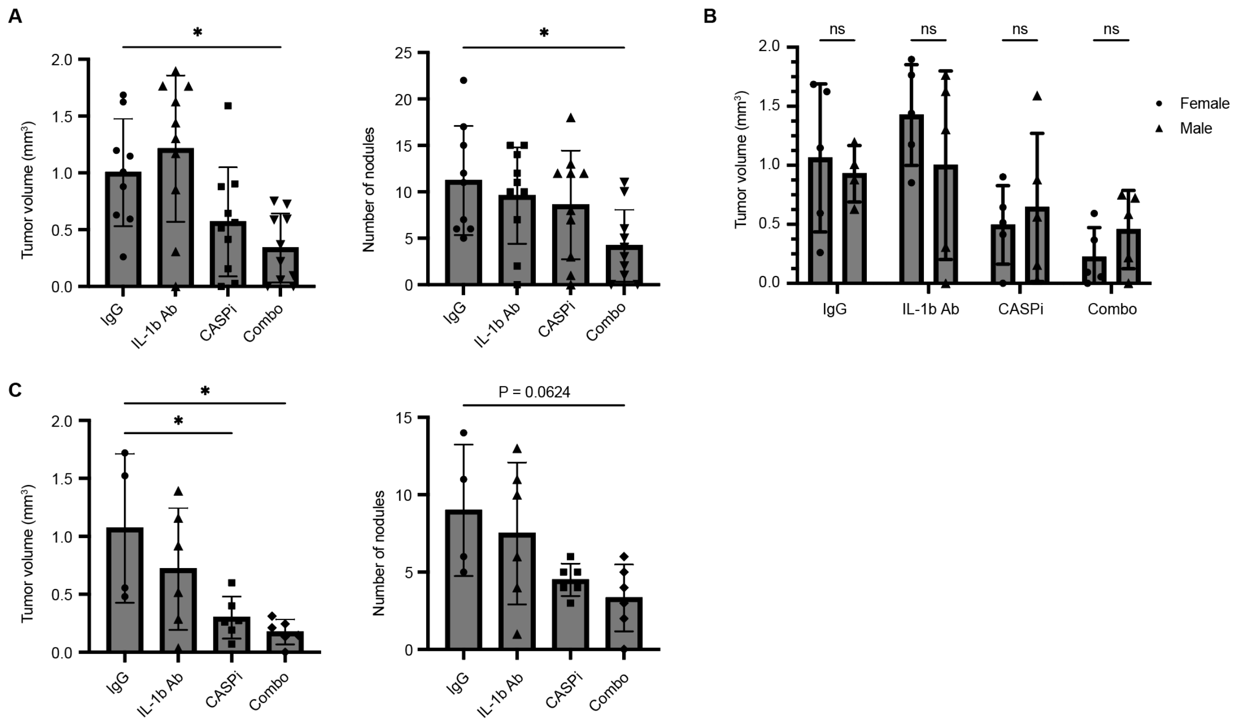


**Figure S11. Treatment of independent KPS cohorts.** (A) Tumor volume quantification and nodule count in first cohort of KPS mice after seven weeks of treatment with IgG, IL-1b antibody, belnacasan (CASPi), and IL-1b antibody and belnacasan (combo). N = 9-10 mice per group. One-way ANOVA with Tukey's multiple comparison test was performed ($p < 0.05$). (B) Separation of first cohort treatment groups by sex shows no significant difference in treatment effect. Two-way ANOVA with Fisher's LSD test was performed ($p < 0.05$). (C) Tumor volume quantification and nodule count of second cohort of KPS mice after seven weeks of treatment. N = 4-6 mice per group. One-way ANOVA with Tukey's multiple comparison test was performed ($p < 0.05$).

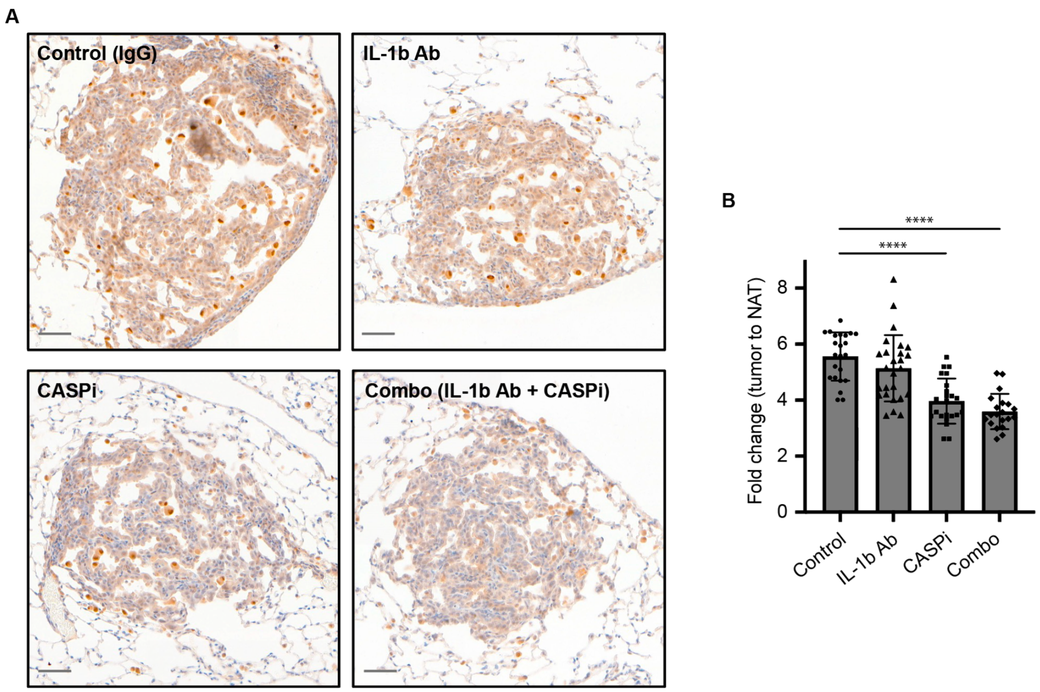


**Figure S12. Immunohistochemical staining of active IL-1b in treated KPS tumors.** (A) Active IL-1b (dark brown) expression was compared across treatment groups. Scale bar 50um. (B) Fold change of mean signal intensity in tumor over NAT was quantified in 2-3 tumors per mouse. N = 2-3 mice per group. One-way ANOVA with Dunnett's multiple comparison test was performed ($p < 0.05$).

**Table S1. Nominated peptide sequences designed to be responsive to interleukin-processing proteases.** Successfully synthesized substrates were used for in vitro recombinant protease FRET screens. Uppercase: L-isomer amino acids; lowercase: D-isomer amino acids; AIBA: α-Aminoisobutyric acid.

| Substrate label | Sequence (C-terminus = CONH2) |
|---|---|
| 1 | GWEHDGG |
| 2 | GWEHDKGG |
| 3 | YVADAP |
| 4 | YVHDAPGG |
| 5 | YVADAPV |
| 6 | GYVADAPVGG |
| 7 | GYVHDAPVGG |
| 8 | GLVCDVGG |
| 9 | GNLLVCDVPG |
| 10 | VDQVDGW |
| 11 | DEVDAP |
| 12 | VDEVDGW |
| 13 | VDQMDGW |
| 14 | EPLFAAR |
| 15 | GPLFAGRGG |
| 16 | GIEPDGG |
| 17 | GIEPDKGG |
| 18 | GIETDGG |
| 19 | IEPDVSQV |
| 20 | GGIVRAGG |
| 21 | GFLG |
| 22 | GGFLGG |
| 23 | GGFGFVGG |
| 24 | GGFAFGG |
| 25 | EPFWEDQ |
| 26 | GEPFWEDQG |
| 27 | GIEPKSDPMPEQ |
| 28 | TPFSGQ |
| 29 | GTPFSGQGG |
| 30 | GAAPLGG |
| 31 | VVLDSEVL |
| 32 | GAYVHGG |
| 33 | GEAYVHGG |
| 34 | VADvRDRQ |
| 35 | GVADvRDRQG |
| 36 | VANvAERQ |
| 37 | GVANvAERQG |
| 38 | VADCADQ |
| 39 | GVADCADQG |
| 40 | GYY[AIBA]NEPGG |
| 41 | VADCRDRQ |
| 42 | VARCADYQ |
| 43 | VADvADYQ |
| 44 | MLDAMGSL |
| 45 | VRRVVQYQD |
| 46 | GDNLLVC GG |
| 47 | APEEIMRRQ |
| 48 | HPVPVYAFSPQ |
| 49 | QPMAVVQSVPQ |
| 50 | GAAPVGG |
| 51 | GAVVASELR |
| 52 | TLLVSGNA |
| 53 | RQRVNWNW |
| 54 | GFIFEEEPIFF G |
| 55 | GEEEPIFGG |
| 56 | GFIFEEEPILCG |
| 57 | GEEEPILGG |
| 58 | GVDVADGG |
| 59 | GLETDGG |
| 60 | GGIETDSG |
| 61 | DGIETDSGVDDD |
| 62 | GELDSGIETDSG |
| 63 | GELQTDGG |
| 64 | GSSIETDALG |
| 65 | VLLVSEVLDD |
| 66 | VVVWSEVVDD |
| 67 | GDRVYIHPFHLD |
| 68 | GCFTERDDD |
| 69 | AAGVAYSEA |

| | |
|---|---|
| 70 | GGVAYSEGG |
| 71 | GALASEIVGG |
| 72 | GARAAEIVGG |
| 73 | GAEAGEIIGG |
| 74 | GASGYGG |
| 75 | GPLGLEEAG |
| 76 | SGVNLDAGFR |
| 77 | DRlVGDDPY |
| 78 | GEGPWLEEEG |
| 79 | GNFDEIDRGG |
| 80 | GRPPGFSPFR |
| 81 | GFIFEEEPGG |
| 82 | GAAYAAYAG |
| 83 | GSAYAATDEGN |
| 84 | GNSAAYASGNG |
| 85 | GPLGLARDDDD |
| 86 | GEPIFFDTWDN |
| 87 | EPILCDSWDDD |
| 88 | GSWDDDDNGG |
| 89 | GLEAIANGG |
| 90 | GLQSITHGG |
| 91 | GIANDGG |
| 92 | GITHDGG |
| 93 | GSAPFSFGG |
| 94 | GSAPYTYGG |
| 95 | GGFAFGISGG |
| 96 | GGSLLGSIQGG |
| 97 | GGSQTDVVGG |
| 98 | GGTVTDVVGG |

| | |
|---|---|
| 99 | GSIITDVVGG |
| 100 | GGHQTDVVGG |
| 101 | GSYATDVVGG |
| 102 | GSIPFDVVGG |
| 103 | GSLTTDVVGG |
| 104 | GSFLTDVVGG |
| 105 | GSYVHDVVGG |
| 106 | GSYVADVVGG |
| 107 | GSWEHDVVGG |
| 108 | GSWFKDVVGG |
| 109 | GSIETDSPGS |
| 110 | GGDETDSPGS |
| 111 | GSVETDSPGS |
| 112 | GSIEPDSLEEGG |
| 113 | GSVEPDSLEEGG |
| 114 | GGDEVDGAVGGG |
| 115 | GGDEVDPAVGGG |
| 116 | GSVIQADGWGG |
| 117 | GRQRRALEKG |
| 118 | GAKIRGQAKG |
| 119 | GSGDRMWggG |
| 120 | GLGALLRVKRLE |
| 121 | GGPQGIWGQ |
| 122 | GGPVPLSLVM |
| 123 | GGfPRSGGG |
| 124 | GGPLGMRGGC |
| 125 | GAIEFDSG |
| 126 | GGYVADAPDGG |

**Table S2. Panel of 20 ABNs for *in vivo* administration.** Each peptide is chemically bonded to a unique mass-coded glutamic acid-labeled fibroprotein and conjugated to 8-arm PEG nanoscaffolds via maleimide-thiol click chemistry. The ABNs were purified with high performance liquid chromatography, and molecular weights were determined with MALDI-TOF. Uppercase: L-isomer amino acids, lowercase: D-isomer amino acids.

| Name | Sequence (N-terminus = NH2, C-terminus = COOH) | MW |
|---|---|---|
| A1 | e(+3G)(+1V)ndneeGFFs(+4A)r-ANP-GGTPFSGQGGC-(PEG8-40kDa) | 64710.9 |
| A2 | e(+2G)Vndnee(+2G)FFs(+4A)r-(ANP)-GGVAYSEGGC-(PEG8-40kDa) | 62528.6 |
| A3 | eG(+6V)ndneeG(+10F)(+10F)sAr-(ANP)-GGAAFAGC-(PEG8-40kDa) | 59565.2 |
| A4 | e(+2G)(+6V)ndneeGFFsAr-(ANP)-GGAIEFDSGC-(PEG8-40kDa) | 59710.9 |
| A5 | eGVndneeGF(+10F)s(+4A)r-(ANP)-GGSGDRMWggGC-(PEG8-40kDa) | 66000.5 |
| A6 | e(+2G)Vndnee(+2G)F(+10F)sAr-(ANP)-GGAKIRGQAKGC-(PEG8-40kDa) | 63514.2 |
| A7 | eG(+6V)ndnee(+3G)(+1F)Fs(+4A)r-(ANP)-GGfPRSGGGC-(PEG8-40kDa) | 60210.3 |
| A8 | e(+2G)(+6V)ndnee(+3G)(+1F)(+1F)s(+1A)r-(ANP)-GGRQRRALEKGC-(PEG8-40kDa) | 64769.9 |
| A9 | e(+3G)(+1V)ndnee(+2G)(+10F)Fs(+4A)r-(ANP)-GGLGALLRVKRLEC-(PEG8-40kDa) | 66952.9 |
| A10 | eGVndnee(+3G)(+1F)Fs(+4A)r-(ANP)-GGYVADAPDGC-(PEG8-40kDa) | 64303.8 |
| A11 | e(+3G)(+1V)ndneeG(+10F)FsAr-(ANP)-GGELDSGIETDSGC-(PEG8-40kDa) | 67842.4 |
| A12 | e(+2G)(+6V)ndneeG(+10F)(+1F)s(+1A)r-(ANP)-GGVDQVDGWGC-(PEG8-40kDa) | 63864.4 |
| A13 | eG(+6V)ndneeG(+10F)Fs(+4A)r-(ANP)-GGPRAAA-Homophe-TSPGC-(PEG8-40kDa) | 65764.8 |
| A14 | e(+2G)(+6V)ndnee(+3G)(+10F)(+1F)s(+4A)r-(ANP)-GGEEKQRIILGC-(PEG8-40kDa) | 66327.0 |
| A15 | eGVndnee(+2G)(+10F)(+10F)s(+4A)r-ANP-GGP-(Cha)-G-Cys(Me)-HAGC-(PEG8-40kDa) | 60949.4 |
| A16 | eG(+6V)ndneeGF(+1F)s(+1A)r-(ANP)-GGRPLGLAGKGC-(PEG8-40kDa) | 63575.6 |
| A17 | e(+2G)Vndnee(+3G)(+10F)(+1F)s(+4A)r-(ANP)-GGPQGIWGQGC-(PEG8-40kDa) | 66158.7 |
| A18 | e(+2G)VndneeG(+10F)(+10F)s(+4A)r-(ANP)-GGPLGMRGGC-(PEG8-40kDa) | 63497.7 |
| A19 | e(+3G)(+1V)ndnee(+2G)(+10F)(+10F)sAr-(ANP)-GGPVGLIGC-(PEG8-40kDa) | 60463.1 |
| A20 | eGVndneeG(+10F)(+10F)sAr-(ANP)-GGPVPLSLVMC-(PEG8-40kDa) | 66457.6 |

**Table S3. Activity zymography probe sequences for *in situ* tissue staining.** Uppercase: L-isomer amino acids; lowercase: D-isomer amino acids; PEG2K: (poly)ethylene-glycol, MW 2000 g/mol; U: succinoyl; X: 6-aminohexanoyl.

| Substrate Name | Sequence (C-terminus = amide) |
|---|---|
| A1 | UeeeeeeeeXGTPFSGQGGrrrrrrrrrX-k(5FAM) |
| A10 | UeeeeeeeeXGGYVADAPDGGrrrrrrrrrX-k(Cy5) |
| A12 | UeeeeeeeeXGVDQVDGWGGrrrrrrrrrX-k(Cy5) |
| polyR | rrrrrrrrrX-k(Cy7) |
